# Protein language models are biased by unequal sequence sampling across the tree of life

**DOI:** 10.1101/2024.03.07.584001

**Authors:** Frances Ding, Jacob Steinhardt

**Affiliations:** Department of Electrical Engineering and Computer Sciences, University of California, Berkeley; Departments of Statistics and Electrical Engineering and Computer Sciences, University of California, Berkeley

## Abstract

Protein language models (pLMs) trained on large protein sequence databases have been used to understand disease and design novel proteins. In design tasks, the likelihood of a protein sequence under a pLM is often used as a proxy for protein fitness, so it is critical to understand what signals likelihoods capture. In this work we find that pLM likelihoods unintentionally encode a species bias: likelihoods of protein sequences from certain species are systematically higher, independent of the protein in question. We quantify this bias and show that it arises in large part because of unequal species representation in popular protein sequence databases. We further show that the bias can be detrimental for some protein design applications, such as enhancing thermostability. These results highlight the importance of understanding and curating pLM training data to mitigate biases and improve protein design capabilities in under-explored parts of sequence space.

## 1 Introduction

Proteins are the building blocks and workhorses of life, performing essential roles in human and ecosystem health. Inspired by advances in natural language processing, many different protein language models (pLMs) have been trained to model the distribution of naturally occurring protein sequences (Rives et al., 2021; Elnaggar et al., 2021; Madani et al., 2023; Lin et al., 2023; Alamdari et al., 2023). pLMs have been successfully used to predict protein 3D structure (Lin et al., 2023), catalytic activity (Eom et al., 2024), and other biophysical properties (Brandes et al., 2022; Jagota et al., 2023), generally with additional supervision for fine-tuning. Excitingly, without needing additional supervision, *likelihoods* from pLMs have been shown to correlate well with protein fitness, i.e. desirable qualities such as catalytic activity, stability, and binding affinity (Meier et al., 2021; Notin et al., 2023a; Nijkamp et al., 2023).

Because of this correlation with fitness, pLM likelihoods are increasingly used in protein design. They have been used to screen for potentially beneficial mutations (Johnson et al., 2023), to design libraries of protein candidates with higher hit rates than previously state-of-the-art synthetic libraries (Shin et al., 2021), and to efficiently evolve human antibodies without any additional supervision (Hie et al., 2023).

In this work we find that likelihoods from popular pLMs have a species bias: likelihoods of naturally occurring protein sequences are systematically higher in certain species, which can be detrimental for some protein design applications. We describe the extent of this species bias, show that it arises from imbalanced species representation in the protein sequence databases used for training, and measure the impact of the bias on protein design.

We first describe a stylized model for pLM training to show intuitively how this bias arises (Section 3), then support our empirical claims with three main results in Sections 4-6. In Section 4 we show that across the many different proteins we study, certain species almost always have higher pLM likelihoods for their protein sequences than other species. For example, in the data we collect, fruit fly proteins have higher likelihoods than the *C. elegans* (roundworm) versions of the same proteins 92% of the time, even though there is no biological reason for fruit fly proteins to be uniformly “fitter” or more canonical. We find consistent species bias in the commonly used Progen2 and ESM2 model families, across several model sizes.

Next, in Section 5 we show that the bias can be largely explained by species representation in protein databases, combined with careful accounting of evolutionary relationships. A handful of species have orders of magnitude more samples than others in the UniProt database (Consortium, 2022) used to train nearly all pLMs. This per-species sample count has a 0.2–0.25 Spearman correlation with the per-species bias, but we find that what really matters is the sample count from *evolutionarily close* species, which achieves a 0.6–0.75 Spearman correlation.

Finally, in Section 6 we examine the implications for protein design. The bias causes protein designs to gravitate towards sequences from favored species. In some cases, this leads the design process to produce worse outcomes. For example, proteins from heat-tolerant microbes are indispensable tools for research and industrial applications because of their stability at high temperatures. However, they are under-represented in sequence databases. Using them as starting points for protein design guided by pLM likelihoods, we find that a majority of designs lose thermostability.

Looking forward, these results suggest that protein designers should use pLM likelihoods carefully and consider whether the species bias should be corrected for a given application. Looking forwards, we believe the protein design field would benefit from more deliberate curation of training data for pLMs, potentially tailored to different applications.

## 2 Related work

**Mismatches between pre-training and downstream tasks** Self-supervised pre-training produces strong results in both natural language processing (NLP) and protein modeling. However, recent work has shown that standard pre-training objectives can imbue language models with properties that are undesirable for downstream tasks, such as lower accuracy when correct outputs contain infrequent words (McCoy et al., 2023), self-delusions (Ortega et al., 2021), and more (Lin et al., 2021). In this work we show how *protein* LMs inherit related properties from their pre-training, and show their impacts on the unique application of protein design.

**Training data bias** Biases in training data are reflected in downstream models. Under-represented subgroups can suffer lower accuracy due to insufficient weight in the training data (Buolamwini & Gebru, 2018; Chen et al., 2018; Kleinberg et al., 2022; Shahbazi et al., 2023), and socially undesirable biases in data are often amplified by models (Bolukbasi et al., 2016; Caliskan et al., 2017; Taori & Hashimoto, 2023). Various papers have studied how re-weighting or curating datasets can mitigate these biases (Zhao et al., 2017; Ryu et al., 2017; Tschandl et al., 2018; Yang et al., 2020), even finding that *overall* performance is improved by over-weighting minority groups and actively increasing diversity in datasets (Gao et al., 2020; Rolf et al., 2021; Lee et al., 2022). In this work we show that despite different training data sampling regimes in popular pLMs, the unbalanced species distribution in the UniProt database (used to train most, if not all, pLMs) creates similar biases across models.

**Protein language models** Many protein language models (pLMs) have been trained using transformers (Rives et al., 2021; Meier et al., 2021; Elnaggar et al., 2021; Hesslow et al., 2022; Ferruz et al., 2022; Elnaggar et al., 2023; Lin et al., 2023; Nijkamp et al., 2023), CNNs (Yang et al., 2022), and other architectures (Alley et al., 2019). pLMs can generate sequences that successfully fold into functional proteins, both by sampling from the model unconditionally, and sampling conditionally on some inputs (Verkuil et al., 2022; Madani et al., 2023; Alamdari et al., 2023). For protein design that seeks to enhance one or more protein fitness properties, pLM likelihoods have been used as a proxy for fitness in the absence of experimental measurements for supervision (Meier et al., 2021; Fannjiang et al., 2022; Hie et al., 2023). If experimental measurements are available, various strategies have been proposed to combine this supervision with pLM outputs to improve fitness prediction (Hsu et al., 2022a; Notin et al., 2023b). Recent work has identified a tendency for pLMs and family-specific protein models to classify a sequence as lower fitness if it has more mutations from a naturally occurring sequence, referred to as sequence similarity bias (Shaw et al., 2023). Our work focuses on a different bias arising from species identity, which we identify specifically in pLMs, and which can compound the effect of sequence similarity bias.

**Figure 1:**
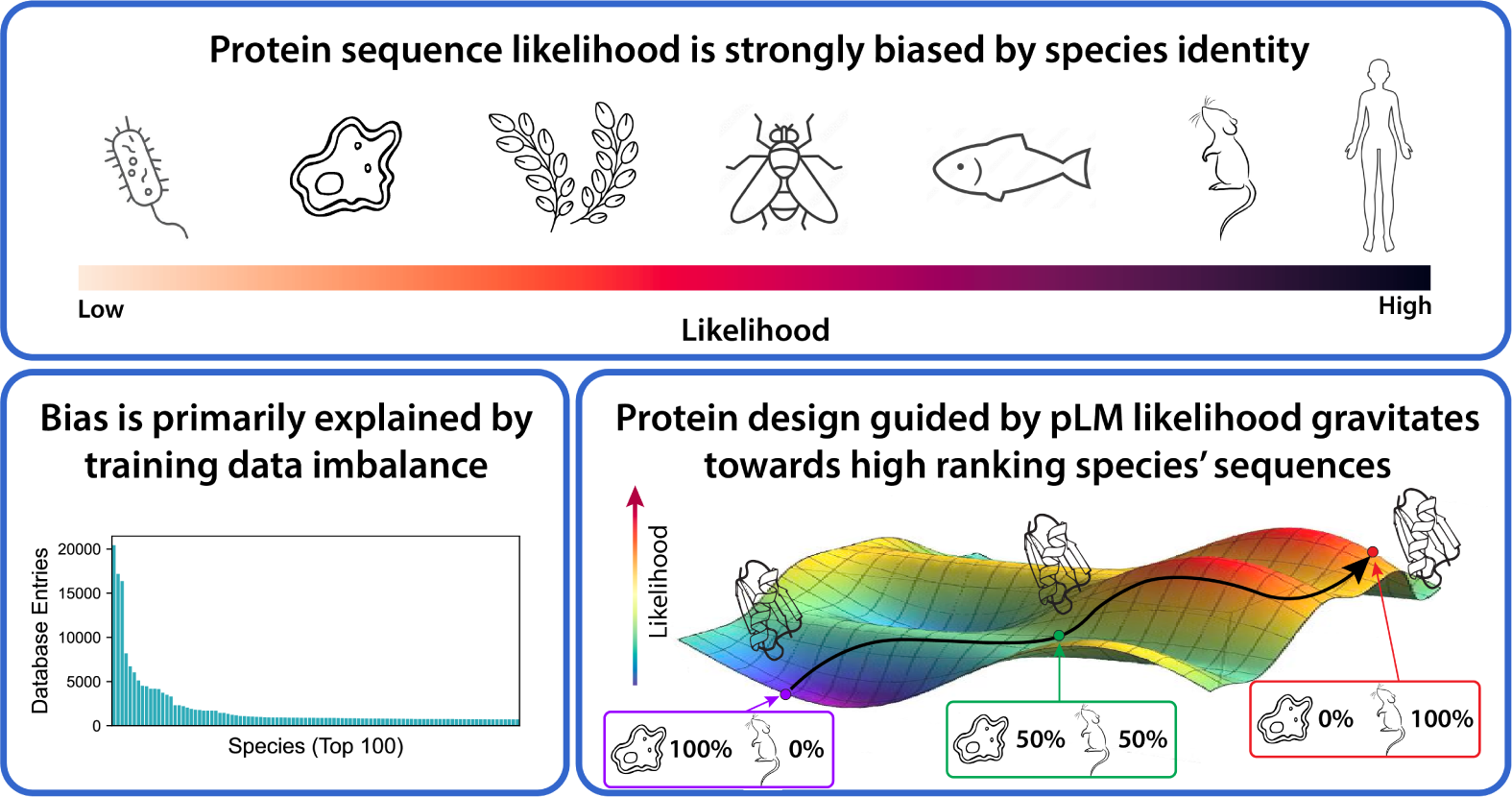
Overview of this paper’s main findings. We find that 1) under popular pLMs, the likelihood of a protein sequence from certain species (e.g. humans, mice, and *E. coli*) is much higher than other species, 2) this bias arises from training data imbalance between different branches of the evolutionary tree of life, and 3) protein design guided by pLM likelihood systematically introduces mutations that increase similarity to high ranking species’ sequences.

## 3 Stylized model for pLM bias

Before presenting empirical results, we provide a brief conceptual example to illustrate why pLMs learn a species bias. pLMs are sequence models with architectures and training objectives inspired by language models trained on text data (Figure 2). Most if not all protein language models are trained on samples from the UniProt protein sequence database, with either an autoregressive (AR) next-token prediction task or a masked language modeling task. For the stylized results in this section, we focus on the autoregressive loss shown below:

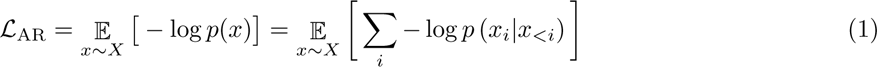

where *x* is a protein sequence, *X* is the distribution of sequences defined by the training set and *i* indexes a single token in *x*.

**Figure 2:**
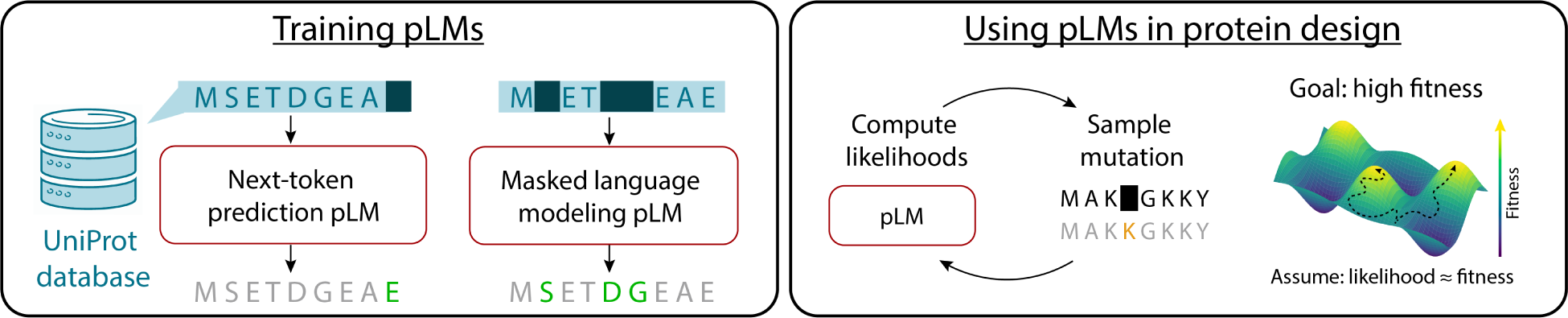
Overview of pLM training and use in protein design. Left: pLMs are trained on amino acid sequences from protein databases (most commonly UniProt) with either next-token prediction or masked language modeling tasks. Right: After training, pLMs may be used directly for protein design by picking a starting sequence and then iteratively generating a set of possible mutations, computing likelihoods of sequences with those mutations, and sampling based on these likelihoods. This process produces designed sequences with high pLM likelihoods, which hopefully corresponds to high fitness.

Consider a stylized model in which protein sequences are represented as vectors *x ∈* R*^D^*, and protein sequences from each species are Gaussian distributed. As a result, the set of all protein sequences forms a mixture of Gaussians.

In this setting, if the training data has *k_s_* samples from species *s*, a pLM that optimizes likelihood (as in Eq. 1) will learn a mixture of Gaussians with weight 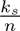.

This stylized setting highlights two trends with likelihoods. First, in the case that different species’ Gaussian distributions have minimal overlap, species that are sampled more, i.e. have larger *k_s_*, will have higher likelihoods (Figure 3a). Second, if some species’ distributions cluster together and overlap significantly, additive effects will make all those species’ sequences have higher likelihoods (Figure 3b).

**Figure 3:**
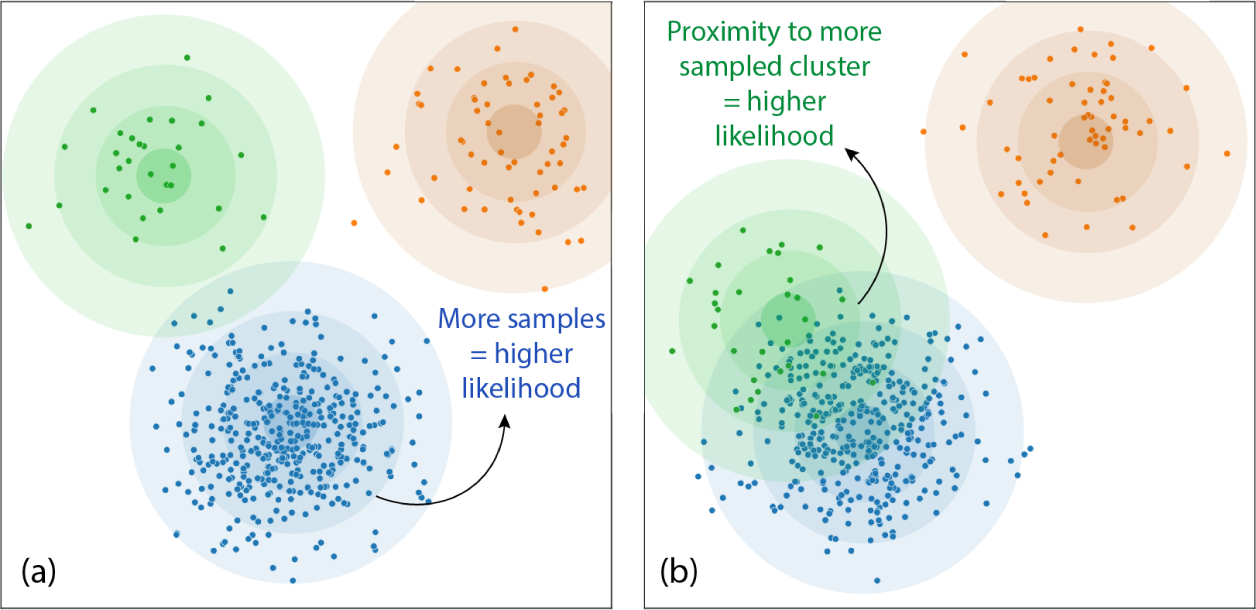
Illustration of biases induced by maximum likelihood training. Left: in well-separated settings, species with the highest prevalence in the training data will have the highest likelihood. Right: when there is overlap between distributions, species clustered together will increase each others’ likelihoods.

We will see both of these effects empirically. In the UniProt database, a few species have far more sequences recorded than others. These species, as well as species in close evolutionary proximity to them, have the highest likelihoods.

## 4 pLM likelihoods are higher for sequences from certain species

We now turn to empirically investigating what factors affect pLM likelihoods. We collect a dataset of orthologous^1^ sequences across the tree of life and across diverse protein types, and we compute pLM likelihoods for each sequence. Unsurprisingly, some protein types have much higher overall likelihoods than others (due to intrinsic disorder, conservation, etc.), but surprisingly, we find that some *species* also have much higher likelihoods than others (across proteins), and that this generalizes across pLMs with different training objectives and data sampling.

**Dataset creation** To create our protein dataset, we started with the top 100 most sequenced species in the UniProt database (Consortium, 2022), filtered for redundancy, then augmented this list with additional model organisms that had whole genomes sequenced, resulting in 133 species total. Next we collected all protein sequences in the Swiss-Prot database (the human-annotated subset of UniProt (Bairoch & Apweiler, 2000)) associated with any of the species in our list. Based on their annotations, we divided the proteins into orthologous sets to be able to compare orthologs to each other. The vast majority of sequences were bacterial, so to create a balanced dataset with many points of comparison between eukaryotes and bacteria, we restricted our attention to proteins with at least 15 eukaryotic orthologs, resulting in 203 distinct protein types, and a total of 7545 sequences in our dataset, 40% being eukaryotic.

**pLMs we study** We focus on two families of pLMs in this work: the Progen2 suite (Nijkamp et al., 2023) (in 5 sizes: xlarge, BFD90, large, base, and medium) and the ESM2 suite (Lin et al., 2023) (in 3 sizes: 15B, 3B, and 650M). These models are among the most popular for downstream use and achieve the best performance among pLMs on many benchmark tasks in ProteinGym (Notin et al., 2023a). Progen2 is an autoregressive transformer trained with next-token prediction on the UniRef90 database (a curated subset of UniProt clustered at 90% sequence identity). ESM2 has a bidirectional transformer architecture and is trained with the masked language modeling objective on data collected in a two-tiered sampling scheme: first randomly select a UniRef50 database member, and then sample a training data point from the UniRef90 cluster that member belongs to. For ESM models, we compute a pseudo-likelihood by masking each token in the sequence, as in Lin et al. (2023).

### 4.1 Results

**Variance explained by species identity** We first investigate what factors explain pLM likelihoods in our dataset. We compute linear regressions of pLM likelihood against species, protein-type, and both at once, and report *R*^2^ values in Table 1. We also compute the fraction of variance explained by the species after controlling for protein type. We find that protein type explains some of the variance, as expected, since proteins vary in prevalence, conservation, and other factors that intuitively affect likelihood. Surprisingly, species identity also explains a significant amount of the variance in pLM likelihoods; for example, for likelihoods from Progen2-xlarge, species accounts for 50% of the variance by itself, and 67% of the variance after controlling for protein type. This suggests that likelihoods have a species bias that holds consistently across the diverse universe of proteins.

**Table 1:** Variance in likelihood explained by species and protein type. 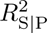 is the fraction of variance explained by species identity, after controlling for protein type^3^.

| MODEL | $R^2_{\text{SPECIES}}$ | $R^2_{\text{PROTEIN}}$ | $R^2_{\text{BOTH}}$ | $R^2_{S\vert P}$ |
| --- | --- | --- | --- | --- |
| PROGEN2-XLARGE | 0.50 | 0.42 | 0.81 | 0.67 |
| PROGEN2-BFD90 | 0.49 | 0.51 | 0.85 | 0.69 |
| PROGEN2-LARGE | 0.46 | 0.60 | 0.87 | 0.67 |
| PROGEN2-BASE | 0.25 | 0.64 | 0.84 | 0.55 |
| PROGEN2-MEDIUM | 0.44 | 0.59 | 0.86 | 0.66 |
| ESM2-15B | 0.25 | 0.42 | 0.60 | 0.32 |
| ESM2-3B | 0.26 | 0.46 | 0.63 | 0.32 |
| ESM2-650M | 0.19 | 0.62 | 0.72 | 0.26 |

We next quantify the bias associated with each species without assuming a linear model of likelihoods. Since each protein is only found in a subset of species, species cannot be fairly compared by a simple average likelihood score. To solve this, we use the Elo rating system, described below, to summarize how often one species has higher likelihoods than another.

**Quantifying species bias via Elo** The Elo rating system was developed to calculate the relative skill levels of players in zero-sum games (Elo, 1978). The difference in two players’ Elo ratings directly translates to the probability of one player winning in a match against the other; for example, the 400 Elo difference between a chess grandmaster and a candidate master implies that the grandmaster is expected to win 90% of matches. In our setting, each time two species have different sequences of the same protein type, we count this as a “match”, where the winner is the species with the higher likelihood for their sequence. If a species has multiple sequences of the same protein, its median likelihood is used to determine the match result. All species start with a baseline Elo rating of 1500, and each pair-wise matchup updates the winner’s rating upwards and the loser’s downwards in a stochastic gradient descent-like step. We use the standard Elo update algorithm with *K* = 32 and average results over 50 permutations of the matchups to ensure results are robust (Boubdir et al., 2023).

Figure 4 plots Elo ratings for each species in our dataset, annotated by phylogenetic taxa. We find that Elo ratings vary widely across species. Using Progen2-xlarge likelihoods, the 25th percentile species (*A. baylyi*) has an Elo rating of 1235 while the 75th percentile species (*S. glossinidius*) has an Elo rating of 1745. This Elo difference of 510 implies that *S. glossinidius* has a higher likelihood for its orthologs 95% of the time. Similarly, if we use ESM2-15B pseudo-likelihoods to compute Elo ratings, there is a 220 Elo difference between the 25th and 75th percentile species, which implies an 80% chance of a higher likelihood. Both models thus have a significant species bias, with Progen2’s being somewhat larger.

**Figure 4:**
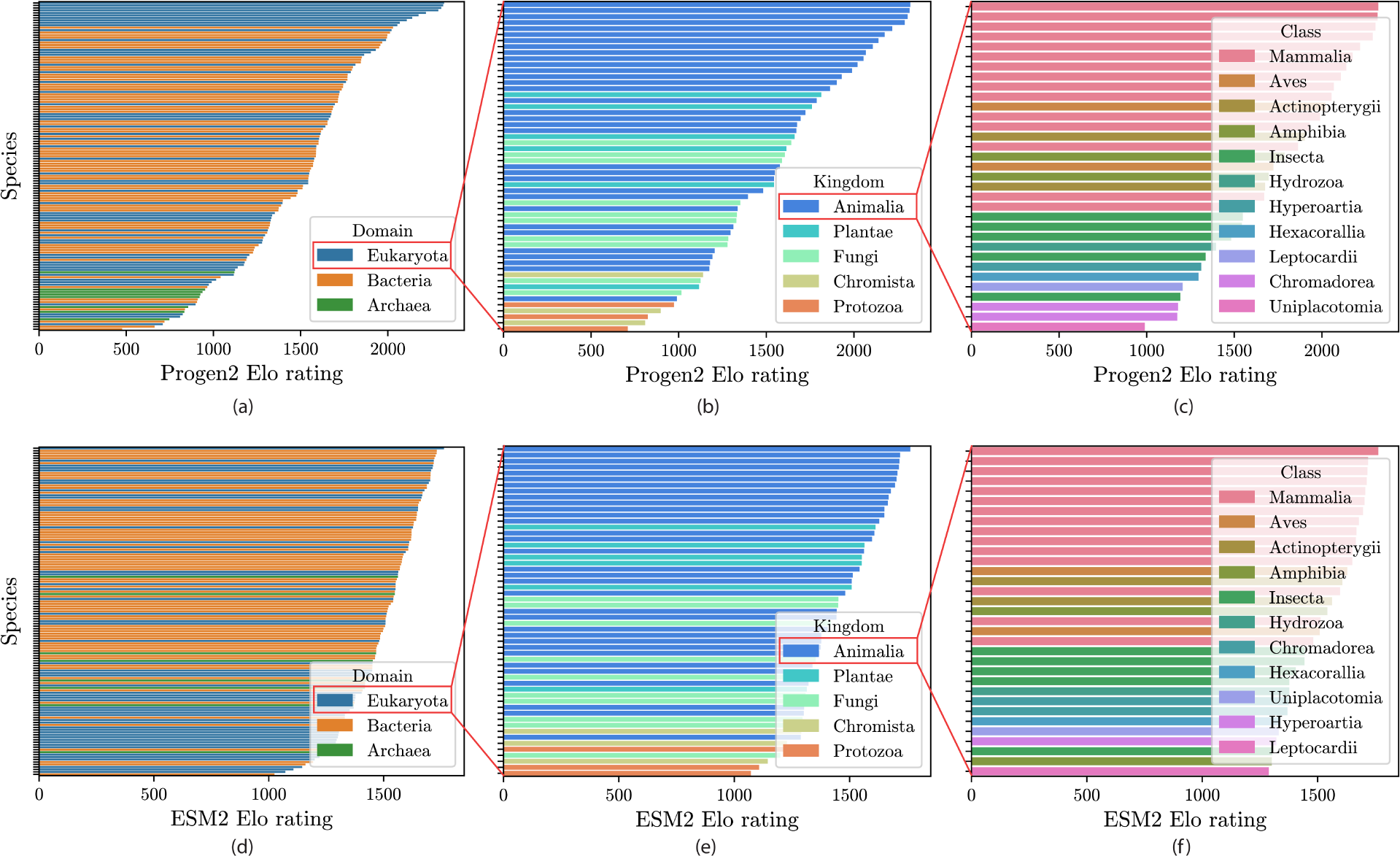
Elo ratings for different species. Elo ratings computed from Progen2-xlarge and ESM2-15B (top and bottom).

Progen2 and ESM2 also show largely similar biases: Elo ratings from Progen2-xlarge and ESM2-15B have a Pearson correlation of 0.83 (see Appendix A for correlations for all pairs of pLMs). Figure 4 further shows that the species bias has some interpretable trends across both models: within eukaryotes, animals have the highest Elo ratings, and within animals, mammals do. This species bias motivates understanding how it arises, which we study in the next section.

## 5 pLM bias is largely explained by species representation in sequence databases

We investigate what factors explain the species bias and find that species representation in popular sequence databases plays a major role. We test an initial hypothesis that Elo ratings will correlate with the number of sequences a species has in a database, and find that this only explains a small part of the bias. As previewed by the stylized model in Section 3, likelihoods may be influenced not only by a given species’ sample counts, but also by evolutionarily-close species’ counts. Taking this second factor into account, we posit a second hypothesis: *Elo ratings will correlate with an “effective” sequence count weighted by evolutionary distance*. We show evidence for this second hypothesis; for example, Progen2-xlarge Elo ratings have a 0.73 Spearman correlation with our effective sequence count metric.

We assess the initial hypothesis by plotting Elo rating against sequence counts in the SwissProt database in Figure 5 (a) and (c). We see that although a few species, such as *H. sapiens*, *M. musculus*, and *E. coli* show the expected trend of high sequence counts and high Elo ratings, most other species are not fit well, with Spearman correlations below 0.3.

**Figure 5:**
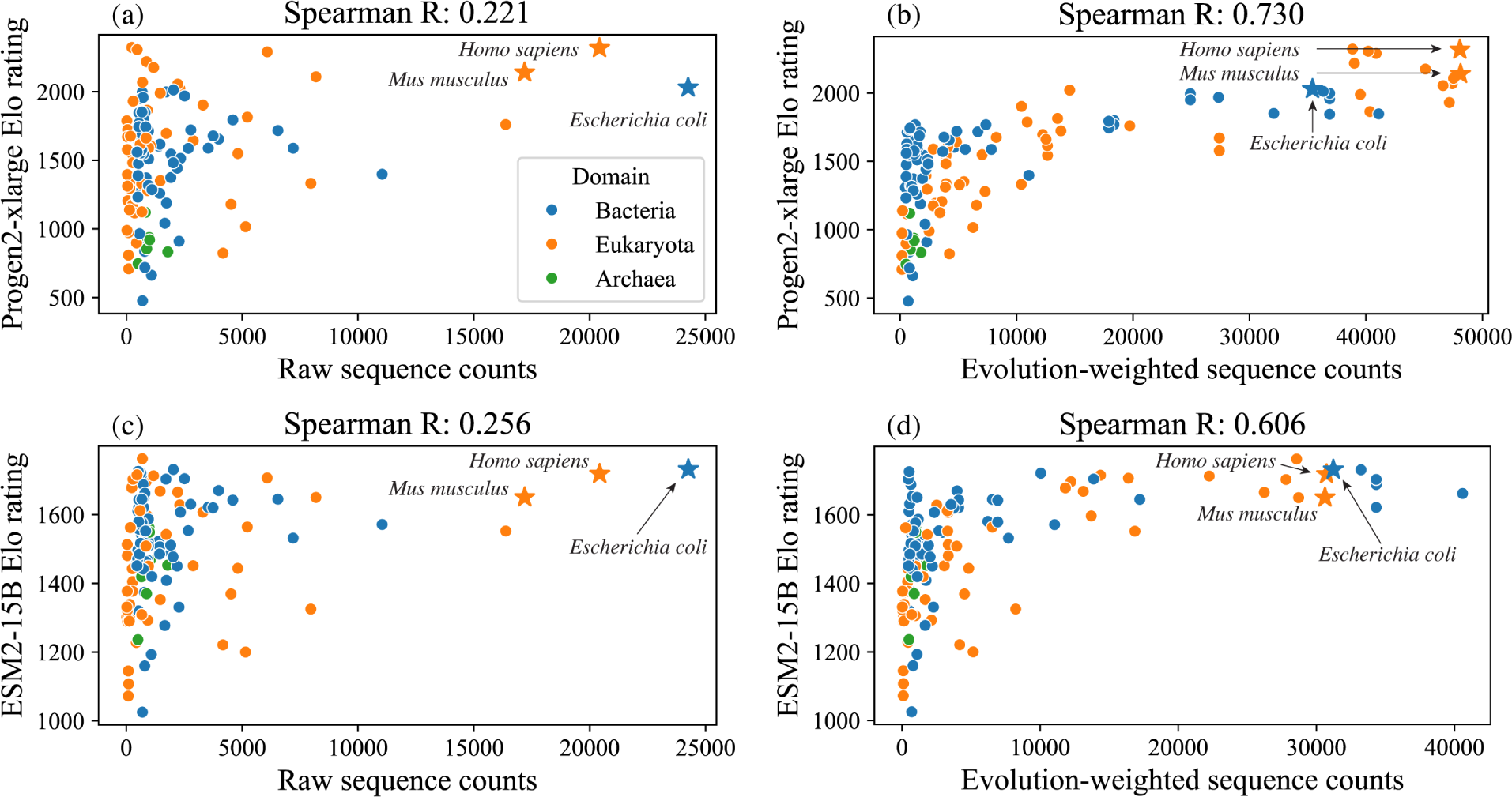
Species Elo ratings plotted against their Swiss-Prot sequence counts and evolution-weighted sequence counts. Top: Elo ratings computed from Progen2-xlarge likelihoods. Bottom: Elo ratings computed from ESM2-15B pseudolikelihoods. Correlation using raw sequence counts (left) is low, while correlation using evolution-weighted sequence counts (right) is high. See Appendix C for similar plots for different choices of database sequence counts such as UniRef50 or UniRef90 counts.

One factor not captured by the initial hypothesis is sequence similarity due to evolution. Figure 6 displays the phylogenetic tree connecting the species in our dataset, annotated with sequence counts and Elo ratings. We see that a few model organisms have extremely large sequence counts, while Elo ratings are spread more diffusely across species. Many species with few sequence counts nonetheless have a high Elo rating, often when the species shares a recent common ancestor with a highly sequenced model organism. We conjecture that the sequence count from a given species contributes to the “effective” sequence count for another species in proportion to the sequence similarity between the two species’ orthologs. Assuming a Poisson model of mutations accumulated over time, sequence similarity between two species’ orthologs is directly related to their evolutionary closeness. Thus we posit our second hypothesis that Elo ratings will correlate with an evolution-weighted sequence count, *n*^evo-weighted^, which we define as follows:

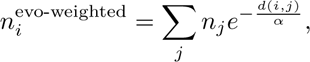

where *n_j_* is the raw sequence count for species *j*, *d*(*i, j*) is the time to last common ancestor between species *i* and *j* collected from the TimeTree of Life resource (Kumar et al., 2022), and *α ∈* R*_≥_*_0_ is a hyperparameter used to scale *d* appropriately. Under the assumption that mutations occur at a fixed rate, 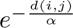 gives the expected overlap in sequence between two species’ orthologs, to approximate the effective sequence counts they contribute to each other^4^.

**Figure 6:**
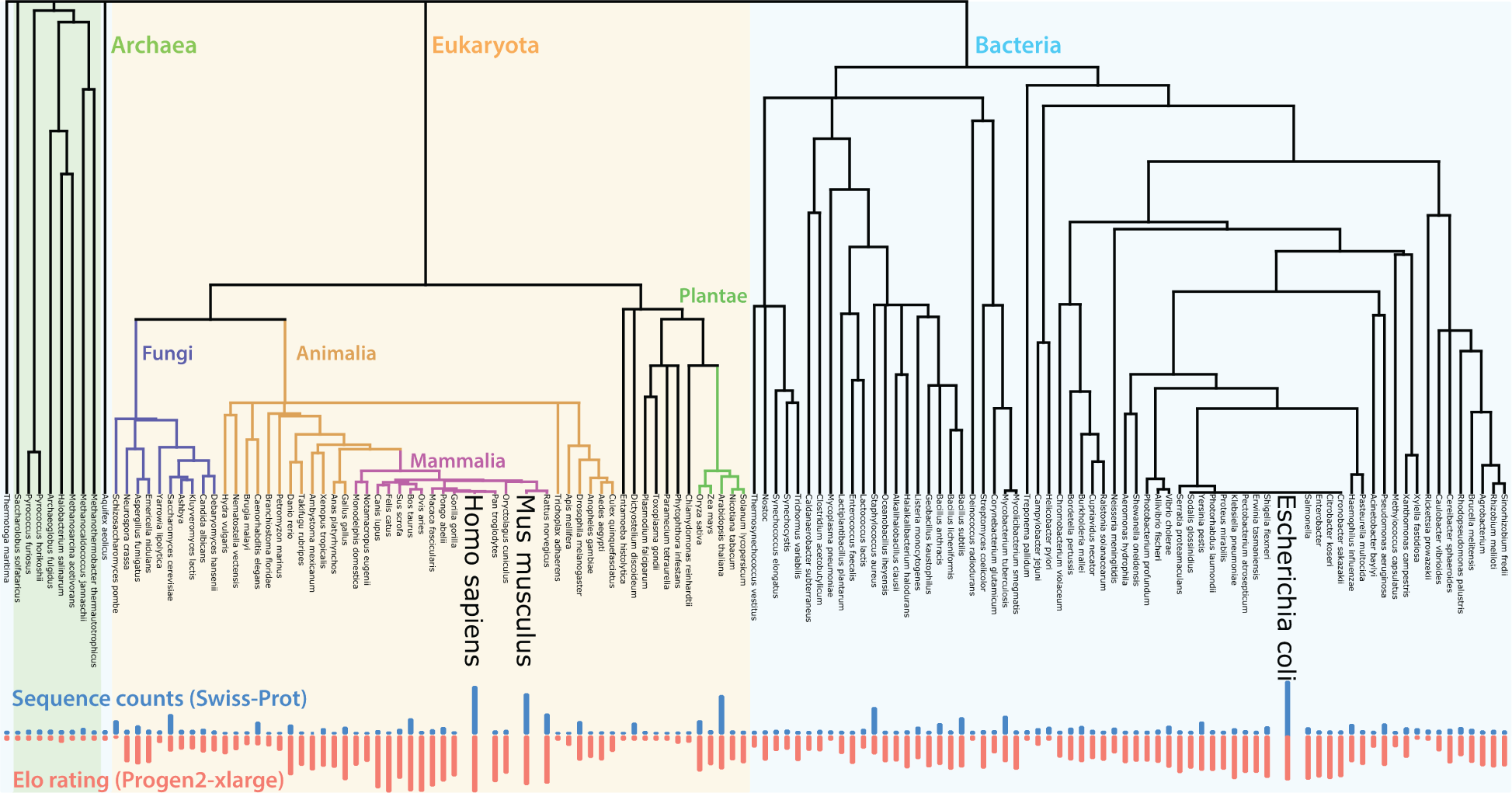
Phylogenetic tree annotated with sequence counts and Elo ratings. The tree’s vertical axis represents time: number of years to the last common ancestor between two species is proportional to the distance from the leaves to the last common node between them.

Figure 5(b, d) plot Elo rating against evolution-weighted sequence counts, and we see that this achieves much higher Spearman correlations than our initial hypothesis: 0.73 and 0.606 for Progen2-xlarge and ESM2-15B Elo ratings, respectively. Thus species representation, with the addition of evolutionary distance, explains a large fraction of the species bias in both pLMs.

It is interesting that both pLMs contain biases with this pattern because their training data differs in important respects. Recall that Progen2 was trained on random samples of UniRef90 (UniProt clustered at 90% sequence identity), while ESM2 used a two-tiered sampling scheme starting with UniRef50 (clustered at 50% identity). One motivation for using UniRef50 clusters is to avoid over-sampling similar, evolutionarily-close sequences. However, our results show that both sampling strategies create a similar species bias, likely because representative sequences for both UniRef50 and UniRef90 clusters are chosen first by annotation quality and second by whether they are from a model organism (Suzek et al., 2015). Additionally, species may have many similar sequences within each cluster, increasing the chances that one of them will be sampled.

## 6 pLM species bias affects protein design

Finally, we investigate the impacts of species bias on protein design with pLMs. Since sequences from high Elo species have higher likelihoods, they may be basins of attraction when designing proteins to optimize likelihood. We test this prediction by simulating a simple protein design workflow and find that designs indeed systematically drift towards sequences from high Elo species.

We then study how this drift affects protein properties. The magnitude of drift is largest when the starting protein is from a low Elo species, and many of the species with the lowest Elo ratings are extremophiles that have adapted to hostile environments such as extreme heat, cold, pH, salinity, and even radiation. Despite their low Elo, these species’ proteins often have unique properties that are useful for a variety of applications, such as research tools, therapeutics, and environmental bioremediation. This suggests that the typical use of likelihood for design will be detrimental when trying to enhance these protein’s properties. We show that this is true for our simulated design workflow. We specifically study thermophilic (heat-loving) and halophilic (salt-loving) species and find that initially thermostable sequences have much lower predicted stability after design, and initially salt-tolerant sequences have lower predicted tolerance after design.

In this section, we first describe our design setup, next show that designs have species drift, and finally show that designs diminish heat and salt tolerance in extremophiles.

### 6.1 Overview of simulated design

In most protein design applications, scientists propose a *set* of candidate designs, with the goal of both high protein fitness and diversity in the set. We follow the design methodology in Zhu et al. (2024) and Fannjiang et al. (2022), who show that optimal tradeoffs between fitness and diversity are achieved by sampling designs from sequence distributions that maximize entropy while satisfying constraints on mean fitness.

Formally, let *X* denote the set of all amino acid sequences of length *L* and let *P* denote set of all distributions over *X*. We aim to sample from the sequence distribution, *p^★^*, that solves the following optimization problem:

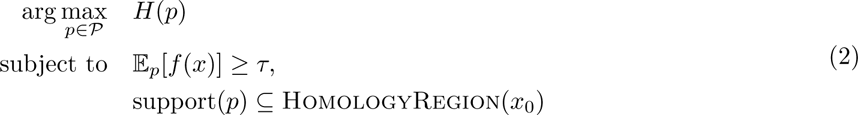

where *H*(*p*) is the entropy of *p*, *f* (*x*) is the sequence log-likelihood under some pLM, *τ ∈* R is a predicted fitness target threshold, and HomologyRegion(*x*_0_) *⊆ X* denotes the region of sequence space that is plausibly homologous to the starting sequence *x*_0_ (to ensure we are designing sequences that still have similar function). Specifically, *x ∈* HomologyRegion(*x*_0_) if and only if the top result returned by a Protein BLAST search (Camacho et al., 2009) for *x* is an ortholog of *x*_0_.

The distribution, *p^★^*, that solves equation 2 has a likelihood of the following form:

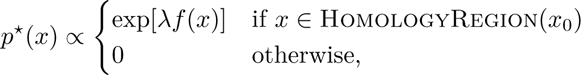

for Lagrange multiplier *λ* which has a one-to-one correspondence with the threshold *τ* in equation 2. Higher *λ* increases the average predicted fitness at the cost of lower diversity.

The distribution *p^★^* is intractable to compute directly, so we use MCMC techniques to sample from it. From the starting sequence *x*_0_, we iteratively propose mutations with Gibbs sampling for 10,000 steps, using Progen2-xlarge likelihoods for the predicted fitness *f* (*x*). We set *λ* = 1 and sample 3 designs with different random seeds for each *x*_0_. In total we collect 1092 designs using many different species’ orthologs as starting sequences, as detailed in the following sections.

### 6.2 Protein design has a species drift

First we show that designs systematically drift towards high Elo species’ orthologs, regardless of starting point. We generate designs for 20 different proteins, each with 15–20 different species’ orthologs as starting points, with the species representing the full range of Elo ratings.

To quantify species drift, we define the *similarity-weighted Elo* of a sequence to reflect the average Elo of species who have orthologs similar to that sequence. Formally:

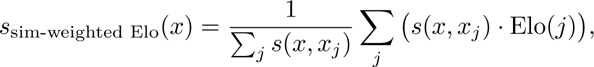

where *x_j_* is the ortholog of *x* from species *j*, *s*(*x, x_j_*) is the sequence similarity between *x* and *x_j_*, and Elo(*j*) is the Elo rating of species *j*. We use *s*(*x, x_j_*) = (1 *−* Levenshtein(*x, x_j_*))^2^, where the Levenshtein distance between two sequences is the minimum number of single-character edits required to change one sequence into another; other choices of *s* lead to similar results (Appendix D).

Figure 7a plots the similarity-weighted Elo of a sequence before and after design. We see that final design sequences tend to increase their similarity-weighted Elo, and we find that this increase is statistically significant under a paired sample *t*-test (*t* = 15.2, *p* = 7.7e-47). This drift holds across the spectrum of Elo ratings and is most prominent for low Elo starting points. This is consequential because many low Elo species are rich sources of useful proteins, as we discuss further in the following section.

**Figure 7:**
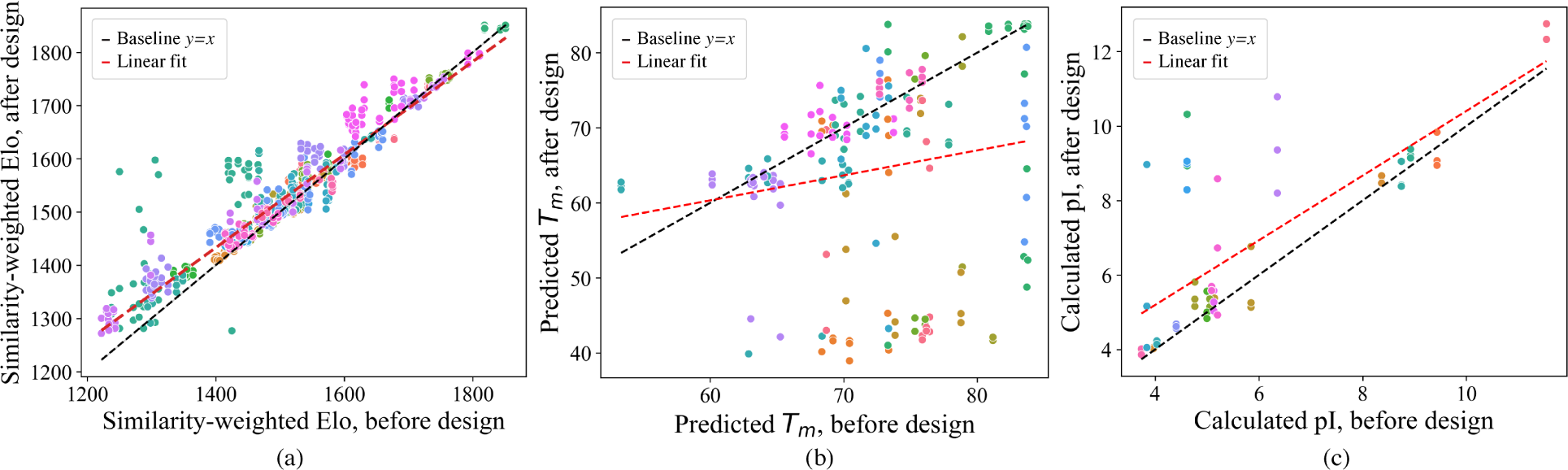
Protein properties before and after protein design. (a) Similarity weighted Elo from design is higher after design. (b) Predicted melting temperature (*T_m_*) is lower (i.e. proteins are less stable) after design. (c) Calculated isoelectric point (pI) is higher (i.e. proteins are less tolerant to salt) after design. Each dot represents one protein design, and color indicates protein type (see Appendix D).

### 6.3 Protein design diminishes extremophile properties

In this section we show that the unique adaptations in extremophile orthologs tend to be diminished after pLM likelihood-based design. Many of the species with the lowest Elo ratings are extremophiles. Despite their low Elo, these species’ proteins are a valuable resource for developing novel biotechnology, with applications including research tools, therapeutics, and environmental bioremediation. For instance, thermophilic (heat-loving) species have evolved proteins with high thermostability (the ability to remain stably folded at high temperatures), which is necessary for many industrial use-cases of engineered proteins (Modarres et al., 2016). Similarly, halophilic (salt-loving) species have evolved proteins to be more acidic (negatively charged) to prevent aggregation in salty environments, and this salt tolerance is crucial to bioremediation strategies such as biofuel production (Daoud & Ben Ali, 2020). Since these thermophilic and halophilic adaptations occur at the individual protein level, they provide perfect test cases to study the effects of protein design.

**Effects on thermostability** To examine protein design’s effects on thermostability, we sample designs from 13 different proteins, each with 3–7 thermophilic species’ orthologs as starting points. We assess the thermostability of each starting sequence and final design *in silico* using the protein melting temperature (*T_m_*) predictor DeepSTABp (Jung et al., 2023).

Figure 7b plots the predicted melting temperature after design vs. before design. We see that designs tend to decrease their predicted melting temperatures, and we find that this decrease is statistically significant under a paired sample *t*-test (*t* = *−*7.1, *p* = 3.5e-11). 63% of designs have lower predicted melting temperatures than their starting sequence, and in one-third of those cases, the predicted decrease is over 20°C. The magnitude of this thermostability decrease is consequential for many applications. For example, engineering strains to successfully ferment bioethanol at 15°C higher temperatures makes a much wider set of raw materials economically feasible for biofuel production (Miah et al., 2022).

**Effects on salt tolerance** To examine protein design’s effects on salt tolerance, we sample designs from 18 different proteins, each with 1–3 halophilic species’ orthologs as starting points. We assess the salt tolerance of a sequence with the isoelectric point (pI) calculator (Kozlowski, 2016), as proteins in halophilic species have lower pI than other species’ orthologs in order to remain stable at high salt concentrations (Gunde-Cimerman et al., 2018).

Figure 7c plots the calculated isoelectric point (pI) after design vs. before design. We see that designs tend to increase their pI, and we find that this increase is statistically significant under a paired sample *t*-test (*t* = 4.6, *p* = 2.0e-5). 83% of designs have higher pI than their starting sequence, and the average pI increase of 4.6 may be consequential–the difference in pH between vinegar and neutral water is approximately 4-5.

Since we assess sequences’ thermostability and salt tolerance *in silico*, these results rely on the accuracy of predictive models. As a sanity check for whether these results would replicate in experimental assays, we analyse the final design sequences and find that a number of them have extremely high sequence similarity (>90%) to mammalian orthologs of the original protein. These orthologs have been experimentally verified to have lower melting temperatures and lower salt tolerance (see Appendix D for more discussion of convergence to orthologs). Thus, even though we only have access to predicted properties, these results suggest that protein design often pushes unique sequences into a basin of attraction around sequences from well-represented species, causing designs to diminish their unique properties.

## 7 Conclusion

In this work we identify and quantify a species bias in pLM likelihoods, trace its origins to uneven sequence sampling across the tree of life, and document its effect of pushing protein designs toward sequences from favored species. This design bias is most likely to be detrimental when the starting point comes from an organism under-represented in sequence databases, and we demonstrate that likelihood-guided design can reduce the thermostability of proteins from heat tolerant species and reduce the salt-tolerance of proteins from species that thrive at high salinity.

It will be important to try to mitigate this species bias with new design algorithms, for example by adjusting the acceptance rate of proposed mutations based on whether the mutation moves towards a common or uncommon ortholog.

However, it is also possible that in some settings the bias will happen to align with design goals. For example, antibody therapeutics are often produced from non-human sources and can generate immunogenic responses in humans (Marks et al., 2021). It would be interesting to test pLM capabilities for humanizing antibodies such that they do not elicit an immune response and thus become safe for therapeutic use.

We focus on likelihoods, but pLM embeddings are also increasingly used in protein design, especially when supervision is available to fine-tune the model. It will be interesting to examine whether embeddings are affected by similar biases and whether they remain after fine-tuning. Future research could also study whether other types of protein models inherit related biases, such as generative models of protein sequences based on 3D structure (Strokach et al., 2020; Hsu et al., 2022b; Dauparas et al., 2022) or generative models of protein structures (Ingraham et al., 2023; Watson et al., 2023). The training data for these models is often even more limited in coverage than the sequence databases used for pLM training.

More broadly, this work highlights the importance of data curation for biological datasets. Training pLMs has only been possible due to decades-long efforts from scientists to standardize sequence information in huge, public databases. While the databases at first primarily served as a resource for protein information queries, today they are additionally treated as defining *distributions* over natural protein sequences. As databases and models continue to grow, it is critical to understand biases present in the data collection process, evaluate whether mitigation of these biases is warranted, and leverage the rich annotations and meta-data in these databases to curate training data with downstream use-cases of models in mind.

## Broader Impacts

Protein design is an active, rapidly changing field of research with potential applications across medicine, biotechnology, and environmental sustainability. Protein language models (pLMs) are one of many computational tools leveraged in protein design efforts, and it is crucial to understand their strengths, weaknesses, and what information they encode. This paper highlights an unintentional bias learned by pLMs that can be detrimental in certain protein design settings. Mitigating this bias may help pLMs become more effective at accelerating protein design, with far-reaching beneficial impacts for human and environmental health.

At the same time, advancements in pLMs, large language models (LLMs), and other biological design tools, such as AlphaFold, have raised the concern that they may be used for the development of biological weapons or other harmful technology. Research based on this paper that improves pLM performance could increase such risks. We hope that by understanding pLMs more thoroughly and collaborating with diverse stakeholders to identify key biosecurity control points, such as physically manufacturing synthetic DNA (Baker & Church, 2024), progress in protein design can achieve its beneficial potential while mitigating risks.

## Acknowledgments

We thank Stephan Allenspach, James Bowden, Moritz Hardt, Milind Jagota, Hanlun Jiang, Erik Jones, Jennifer Listgarten, Thi Nguyen, Hunter Nisonoff, Alex Pan, Jeremy Reiter, Yun Song, and Junhao Xiong for helpful discussions. FD is supported by the NSF Graduate Research Fellowship Program under Grant No. DGE 1752814 and the Open Philanthropy AI Fellowship Program. JS is supported by the NSF SaTC CORE Award No. 1804794 and the Simons Foundation.

## Footnotes

1 Orthologs are genes/proteins in different species that evolved from a common ancestral gene/protein by speciation, and, in general, orthologs retain the same function during the course of evolution.

3 Example 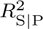 derivation: 0.67 = (0.81 - 0.42)/(1 - 0.42)

4 Mutation rates in fact vary across species and proteins, so as more species’ proteomes are annotated, one could compute a more precise effective sequence count by directly calculating the average sequence similarity between each species’ proteomes, and using this in place of the 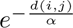 term.

### A Elo ratings from different pLMs

Figure 8 plots the correlation between Elo ratings computed from different pLMs. We see that pLMs within the same family have nearly identical Elo ratings, and pLMs across families also correlate highly.

**Figure 8:**
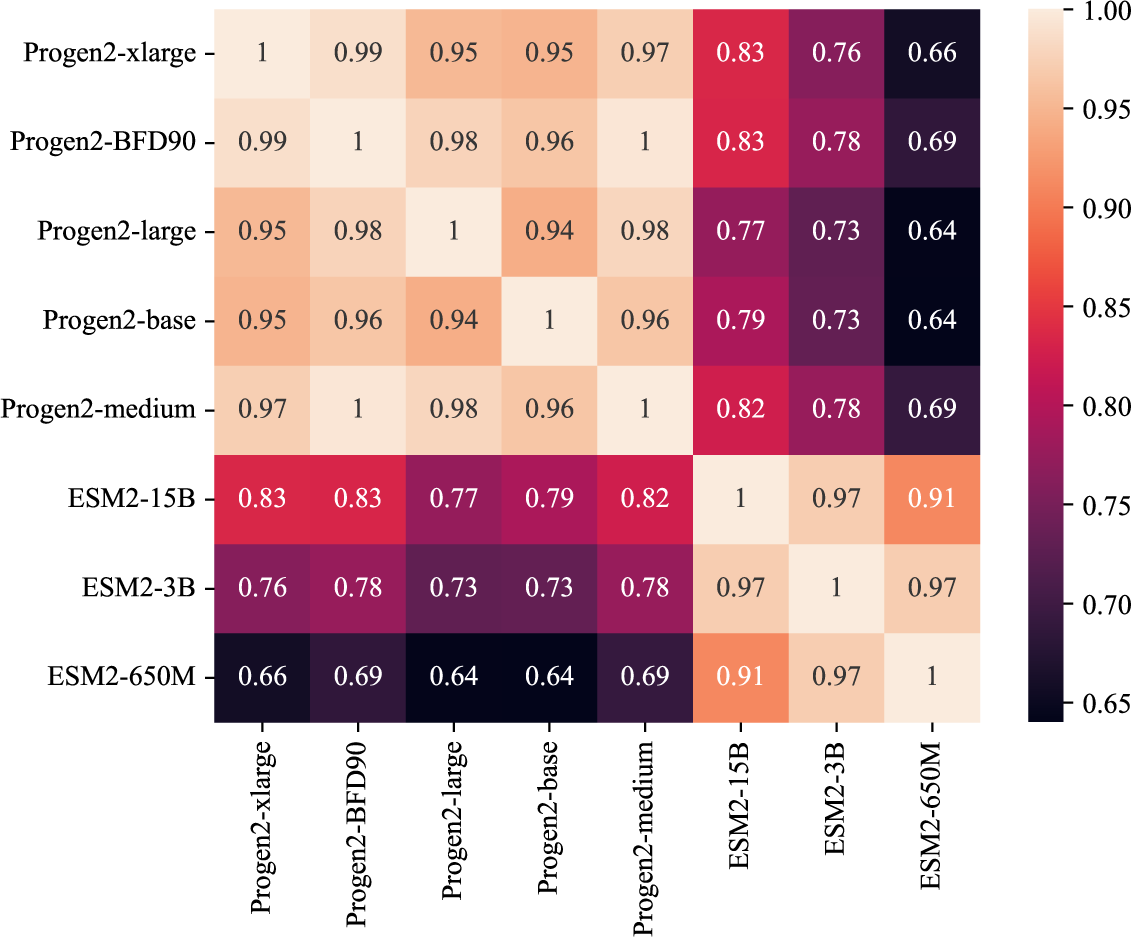
Heatmap of the Pearson correlation between Elo scores from different pLMs.

### B Dataset creation details

We started with the most represented species in the Swiss-Prot database (available from the UniProt statistics page) filtered for redundancy, then added organisms with annotated genomes from the NCBI Genome Server to include at least one organism from each listed category (e.g. Fish, Insects). We also required each organism to have at least 100 entries in the Swiss-Prot database, which resulted in 133 final species. We next used the UniProt REST API to download all Swiss-Prot sequences from each of the selected species. We divided the proteins into orthologous sets based on the protein name and function annotations. The majority of sequences collected were bacterial, so to create a more balanced dataset that allowed for comparison between eukaryotes and bacteria, we kept a protein in our dataset if they had at least 15 eukaryotic orthologs.

### C Robustness to different database sequence counts

In the main text, Figure 5 shows the correlation between Elo scores and raw sequence counts, and between Elo scores and evolution-weighted sequence counts, using the number of SwissProt entries per species as the raw sequence count. Here we show that we find similar results by using other choices of raw sequence counts. Figure 9 shows results with UniRef90 sequence counts. Figure 10 shows results with sequence counts tallied from two-tiered sampling from UniProt: first sample a representative member of UniRef50, then sample a protein sequence from the UniRef90 cluster that the member belongs to. In all cases, correlation with Elo ratings is significantly higher with evolution-weighted sequence counts.

**Figure 9:**
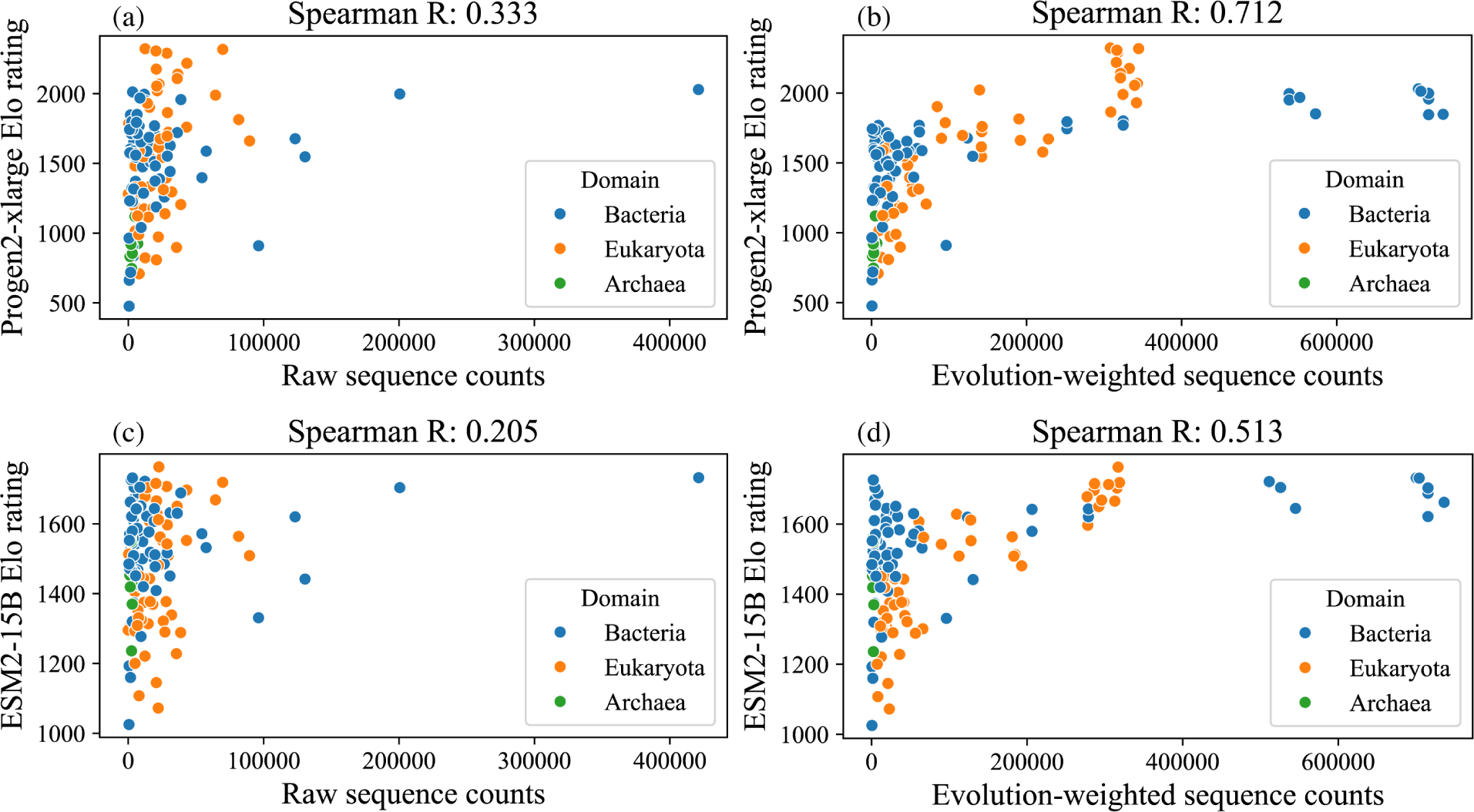
Species Elo ratings plotted against their UniRef90 sequence counts and evolution-weighted sequence counts. Top: Elo ratings computed from Progen2-xlarge likelihoods. Bottom: Elo ratings computed from ESM2-15B pseudolikelihoods. Correlation using raw sequence counts (left) is low, while correlation using evolution-weighted sequence counts (right) is higher.

**Figure 10:**
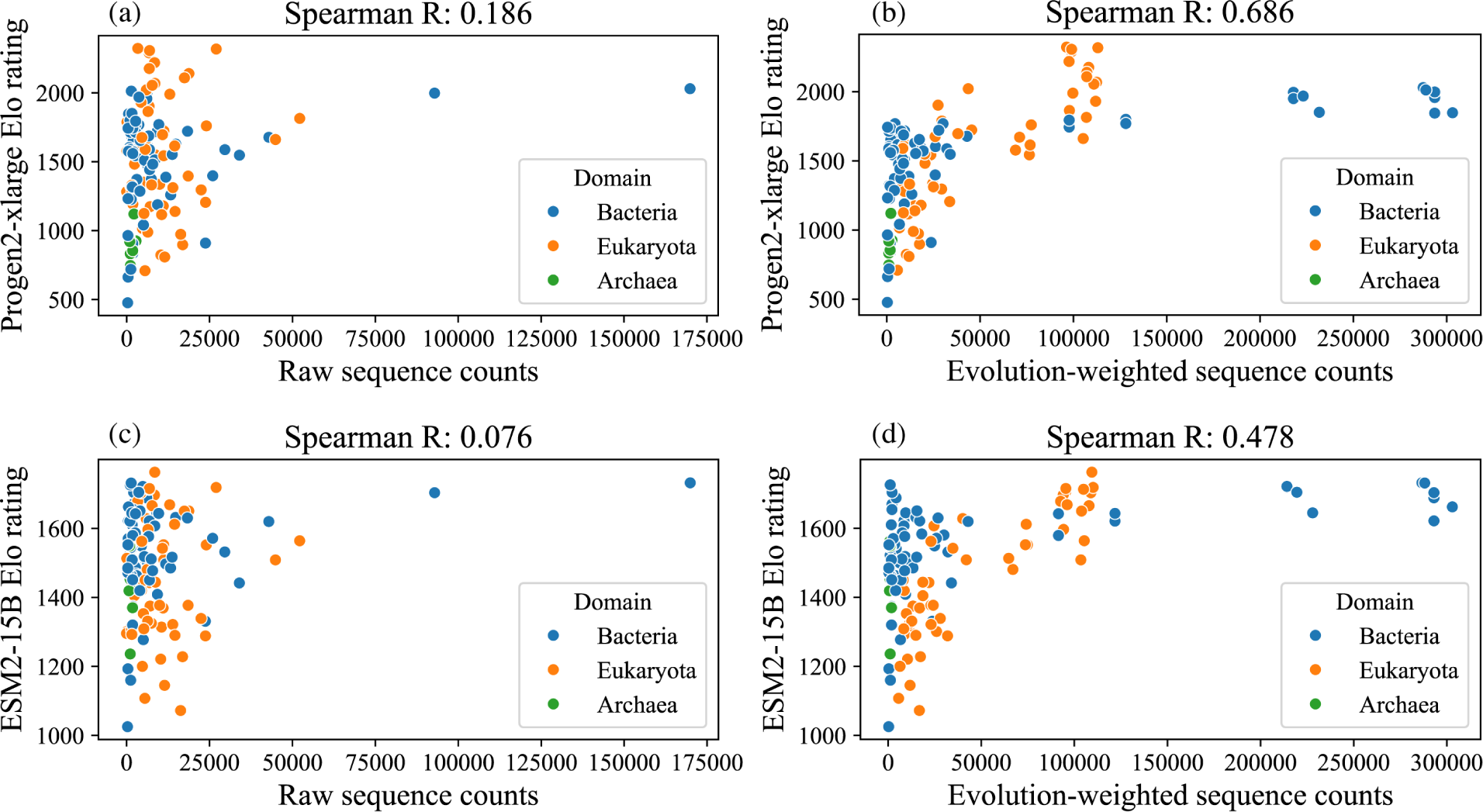
Species Elo ratings plotted against sequence counts from two-tiered sampling: first sample a representative from UniRef50 and then sample a sequence from the UniRef90 cluster of the representative. Top: Elo ratings computed from Progen2-xlarge likelihoods. Bottom: Elo ratings computed from ESM2-15B pseudolikelihoods. Correlation using raw sequence counts (left) is low, while correlation using evolution-weighted sequence counts (right) is higher.

### D Details on protein design experiments

#### D.1 Details for similarity-weighted Elo

For the results in the main text, we compute similarity-weighted Elo with *s*(*x, x_j_*) = (1 *−* Levenshtein(*x, x_j_*))^2^. Here we show that results are robust to other choices of *s*. We use the Bio.Align package to score the optimal alignment between sequences with the Smith-Waterman algorithm, using the BLOSUM62 matrix to score the penalties for each amino acid substitution. Similar amino acids receive a smaller edit penalty compared to more chemically distinct amino acids, and we use the maximum alignment score as the similarity *s*. In contrast, the Levenshtein distance assigns an equal penalty to all substitutions. Figure 11 plots similarity-weighted Elo after design vs. before design. We see that for both similarity metrics, similarity-weighted Elo increases after design.

**Figure 11:**
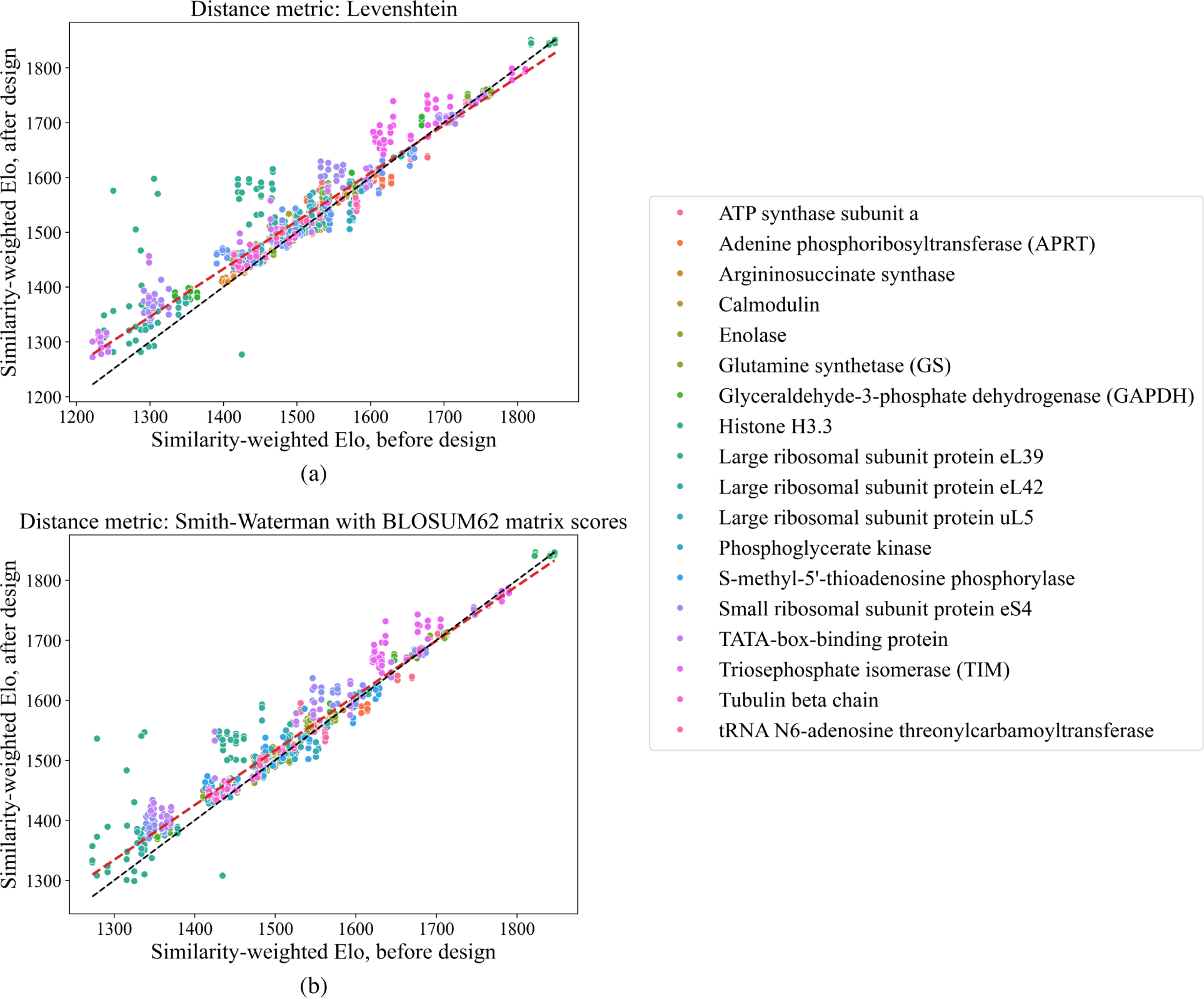
Similarity-weighted Elo before and after design. Top: similarity-weighted Elo computed using Levenshtein distance. Bottom: similarity-weighted Elo computed using the Smith-Waterman algorithm to compute a local alignment under BLOSUM62 matrix scores for amino acid substitutions.

Proteins were selected for design only if they had at least 15 orthologs in Swiss-Prot and if the Elo ratings of the species with orthologs spanned at least 200. Under the computational constraints of how many designs could be generated, proteins were chosen to cover different types of functions (e.g. enzymes, structural proteins, etc.)

#### D.2 Details for thermostability

We study the 7 thermophilic species out of the original 133 species collected in our dataset: *Methanocaldococcus jannaschii*, *Archaeoglobus fulgidus*, *Methanothermobacter thermautotrophicus*, *Methanosarcina acetivorans*, *Pyrococcus furiosus*, *Pyrococcus horikoshii*, and *Saccharolobus solfataricus*. Figure 12 plots the same data as Figure 7b with the legend added. Proteins were selected for design only if they had at least 3 orthologs from thermophilic species, and under the computational constraints of how many designs could be generated, proteins were chosen to cover different types of functions (e.g. enzymes, structural proteins, etc.)

**Figure 12:**
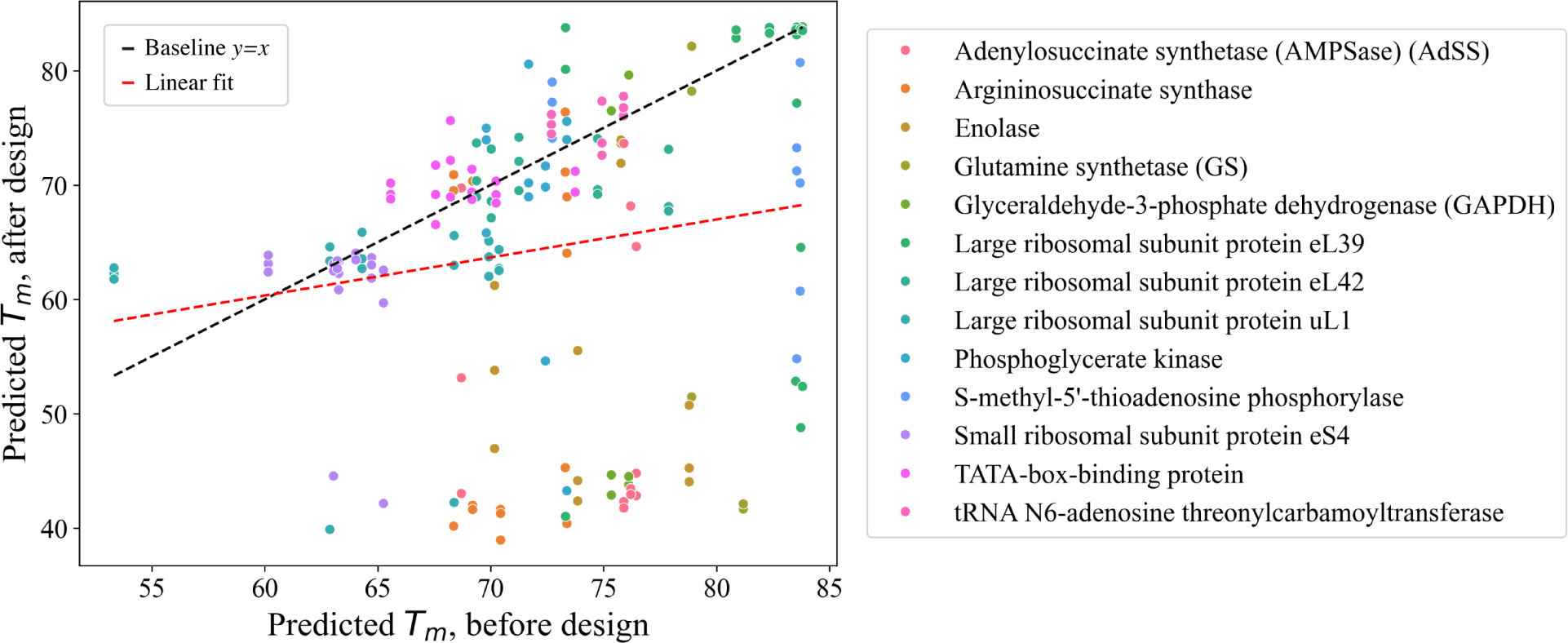
Predicted melting temperature (*T_m_*) after design vs. before design.

#### D.3 Details for salt tolerance

We study the 3 halophilic species out of the original 133 species collected in our dataset: *Halobacterium salinarum*, *Alkalihalobacillus clausii*, and *Halalkalibacterium halodurans*. Figure 13 plots the same data as Figure 7c with the legend added. The proteins used for design in this experiment are all ribosomal because ribosomal proteins are generally basic, and thus provide a focused test case to study whether halophilic species’ orthologs become more basic after design.

**Figure 13:**
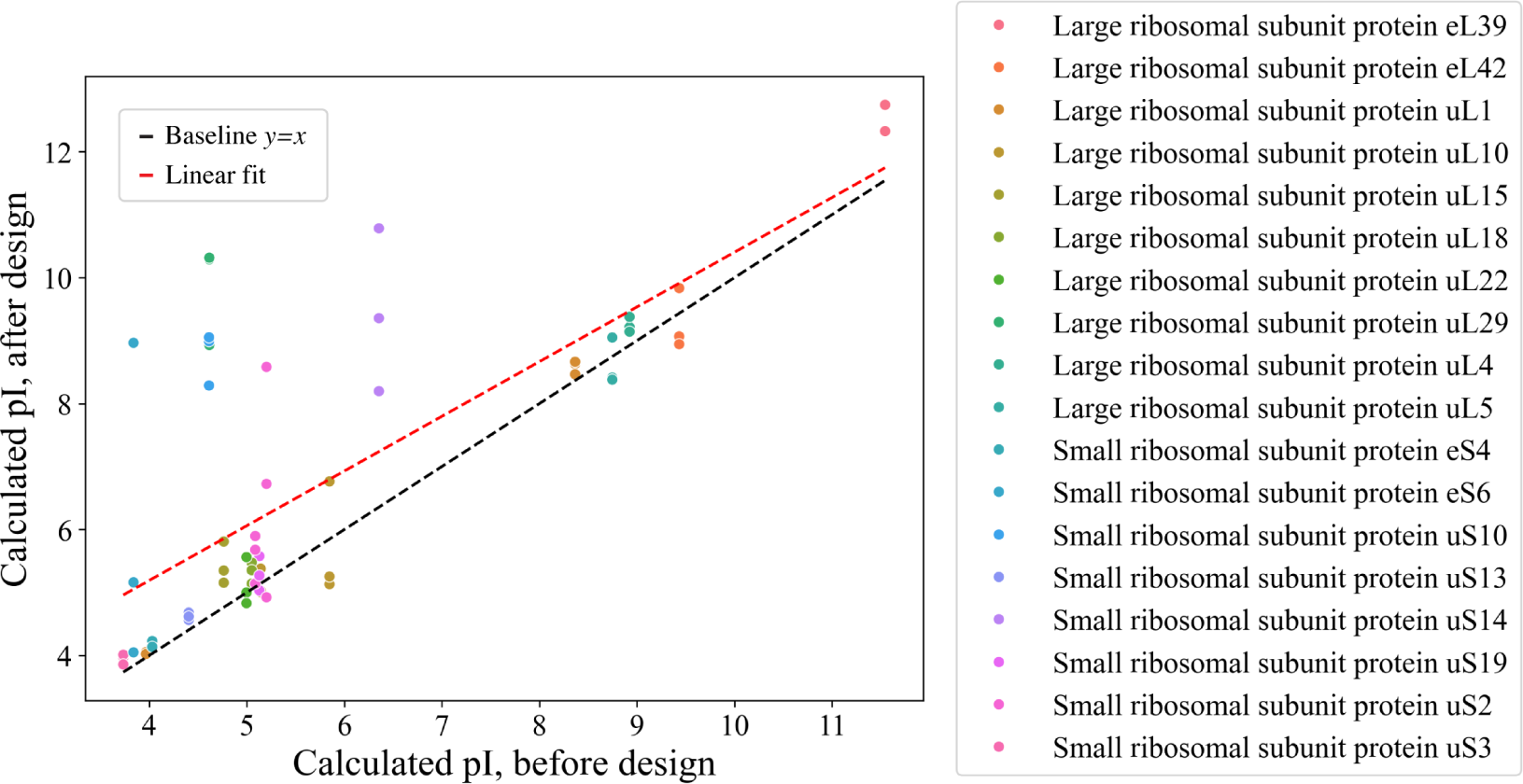
Calculated isoelectric point (pI) after design vs. before design.

#### D.4 Convergence to high Elo orthologs after design

We find that some protein designs result in sequences nearly identical to orthologs from high Elo species. To quantify this, we compute the fraction of designs that “converge” to a ortholog in our dataset, when we set the threshold for convergence to be 90%, 95%, and 98% sequence identity. Figure 14 plots these convergence frequencies. We see that the convergence rate varies significantly between different proteins, and that convergence happens much more often to a higher Elo species’ ortholog compared to a lower Elo species’ ortholog.

**Figure 14:**
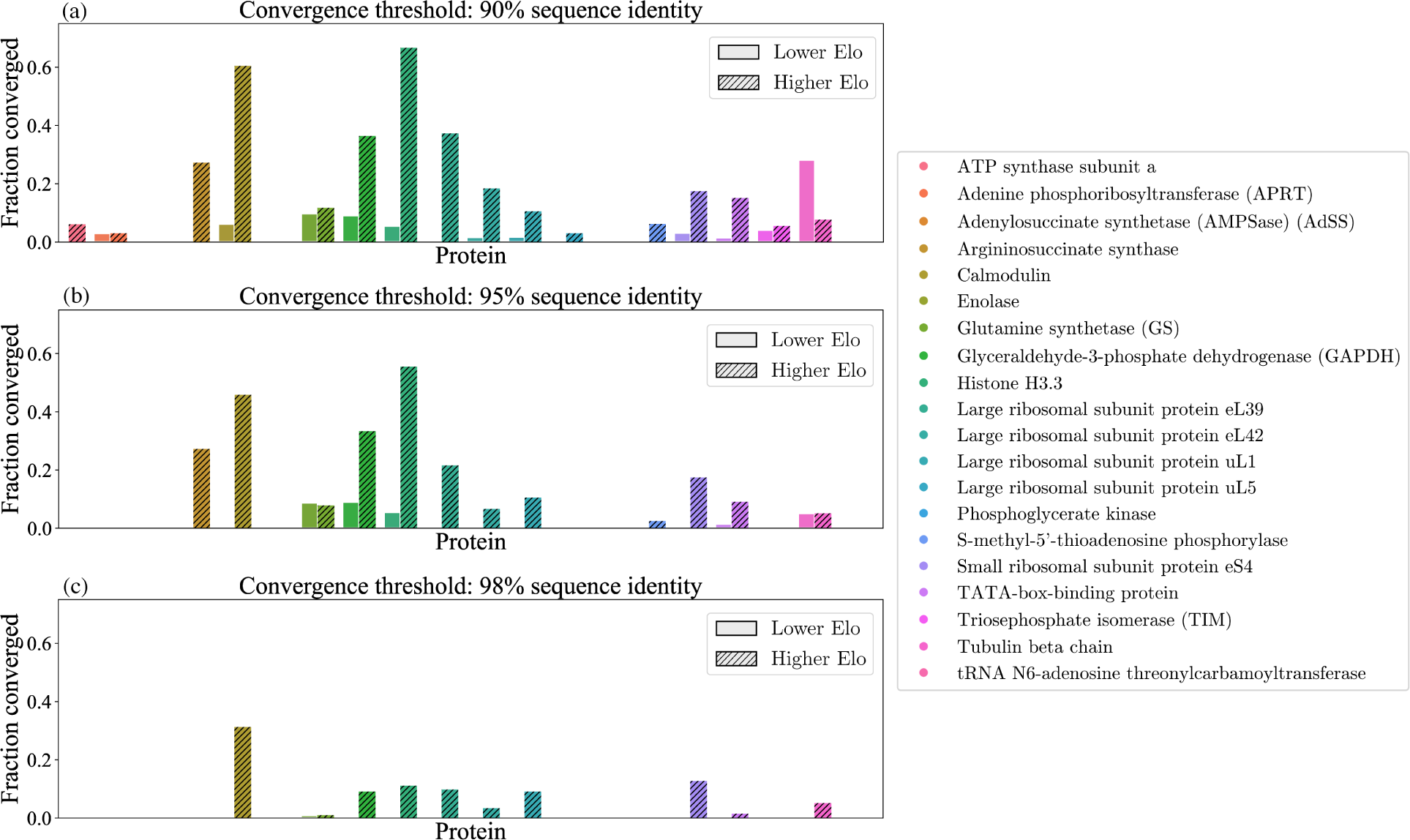
Frequency of convergence to naturally-occurring orthologs from a different species. Convergence to a lower Elo species’ ortholog is represented by the solid bars, and convergence to a higher Elo species’ ortholog is represented by the shaded bars. The threshold for convergence is 90% sequence identity, 95% sequence identity, and 98% sequence identity for the top, middle, and bottom, respectively.

